# Modulation of PKM1/2 levels by steric blocking morpholinos alters the metabolic and pluripotent state of murine pluripotent stem cells

**DOI:** 10.1101/2021.11.23.469697

**Authors:** Joshua G. Dierolf, Hailey L.M. Hunter, Andrew J. Watson, Dean H. Betts

**Affiliations:** Department of Physiology and Pharmacology, Schulich School of Medicine & Dentistry, The University of Western Ontario, London, Canada; Department of Obstetrics and Gynecology, Schulich School of Medicine & Dentistry, The University of Western Ontario, London, Canada.; The Children’s Health Research Institute (CHRI), Lawson Health Research Institute, London, Canada.

**Keywords:** embryonic, formative, metabolism, morpholino, naïve, pluripotency, primed

## Abstract

Cellular metabolism plays both an active and passive role in embryonic development, pluripotency, and cell-fate decisions. However, little is known regarding the role of metabolism in regulating the recently described “formative” pluripotent state. The pluripotent developmental continuum features a metabolic switch from a bivalent metabolism (both glycolysis and oxidative phosphorylation) in naïve cells, to predominantly glycolysis in primed cells. We investigated the role of pyruvate kinase muscle isoforms (PKM1/2) in naïve, formative, and primed mouse embryonic stem cells through modulation of PKM1/2 mRNA transcripts using steric blocking morpholinos that downregulate PKM2 and upregulate PKM1. We have examined these effects in naïve, formative, and primed cells by quantifying the effects of PKM1/2 modulation on pluripotent and metabolic transcripts and by measuring shifts in the population frequencies of cells expressing naïve and primed cell surface markers by flow cytometry. Our results demonstrate that modulating PKM1 and PKM2 levels alters the transition from the naïve state into a primed pluripotent state by enhancing the proportion of the affected cells seen in the “formative” state. Therefore, we conclude that PKM1/2 actively contributes to mechanisms that oversee early stem pluripotency and their progression towards a primed pluripotent state.

## Introduction

Pluripotent stem cells (PSCs) are characterized by their unlimited self-renewal and potential to specialize into all cell types of the adult organism. Approximately 3.5 days following fertilization (E3.5), the mouse embryo contains a niche of cells within the blastocyst called the inner cell mass (ICM) (1,2). This cell niche represents the earliest pluripotent stem cell (PSC) population of the developing embryo and are the origin of the primary germ layers that result in the formation of the fetus. ICM cells also represent the origins of mouse embryonic stem cells (mESCs) that are important research models for unraveling early developmental cell fate control mechanisms and are important resources for developing potential cell-based therapeutics. Mouse ESCs can be explanted from the embryo until E8.0, however, several key differences reflecting changes in their pluripotency status arise between the E3.5 and E8.0 (3). Some of these differences include developmentally programmed changes in gene expression, epigenetic landscape, metabolic preferences, and capacity to contribute to all germ cell layers and foster chimeric development (4,5). Explanted mESCs between E3.5 and E4.5 and mouse epiblast stem cells (mEpiSCs) from epiblasts between E7.25 and E8.0 both express core pluripotency genes including sex determining region Y – box 2 (Sox2), octamer-binding transcription factor 4 (Oct4), and Nanog (lower in mouse epiblast stem cells; mEpiSCs). However, mESCs express naïve pluripotency associated genes such as reduced expression 1 (Rex1), platelet endothelial cell adhesion molecule 1 (Pecam-1), and orphan nuclear receptor Esrrb at greater levels than mEpiSCs (6). Conversely, mEpiSCs express primed pluripotency genes such as Zinc finger protein 2 (Zic2), T(Brachyury), and Cerberus (Cer1) more so than mESCs (3). Recent studies have indicated there is an intermediate state that exists between the naïve and primed ends of the pluripotency spectrum referred to as ‘formative pluripotency’, representative of the E5.5-E6.0 post-implantation epiblast (7). Therefore, E3.5-E4.5, E5.5-6.0, and E7.25-E8.0 represent distinct states of the pluripotency continuum, and are referred to as naïve, formative, and primed pluripotent states respectively, and represent the ICM cells of the pre- and post-implantation epiblast stages (3,4,8,9).

This newly defined formative pluripotent state is consistent with the phased progression model suggesting that all differentiating naïve cells must phase through the formative, then primed state before exiting pluripotency(7). The formative state cells are precursors to primordial germ cells. Unlike mESCs or mEpiSCs, formative state stem cells respond directly to germ cell induction and colonize the germline in chimeras (9-11). Ground state mESCs that are chemically transitioned towards a primed pluripotent state through the replacement of LIF/2i with activin and fibroblast growth factor (Fgf) supplementation (FA), hereafter referred to as mouse epiblast-like cells (mEpiLCs), do not fully commit to primed pluripotency and exhibit an intermediate potency with the potential to differentiate into mPGCLCs (10,12). Formative state mEpiLCs show increased expression of genes including *de novo* DNA methyltransferase 3a/b (Dnmt3a/b), fibroblast growth factor 5 (Fgf5), Sal-like protein 2 (Sall2), Sox3, and POU domain class 3 transcription factor 1 (Oct6; Pou3f1), following a decrease in Nanog expression (7,13). The switch from metabolic bivalency to aerobic glycolysis also occurs during this transitional phase (14).

There is growing evidence showing that not only are the metabolic preferences of naïve and primed pluripotent states distinct, but their metabolic preferences also promote developmental processes, maintain their pluripotency state, and prime cell fate decisions (15-17). The concept of metabolic remodelling and reprogramming has been demonstrated in a variety of stem cells including T cell fate control, direct reprogramming of glial cells to neurons, neuronal metabolic preferences during differentiation, and improving stemness through mitochondrial function by nicotinamide adenine dinucleotide (NAD) reduction (18-21). On either end of the pluripotent continuum, naïve and primed states observe unique preferential metabolic phenotypes. These *in vitro* metabolic preferences may exist as a by-product of their *in vivo* correlate’s metabolism due to restricted, otherwise hypoxic, physiological oxygen levels that the pre- and early post-implantation embryo develops within. The recently described and stabilized intermediate pluripotent state, the formative state, has yet to be metabolically profiled, however, as this interval is representative of the early post-implantation blastocyst, it could demonstrate a bias for aerobic glycolysis (7,9).

Pyruvate kinase muscle isoforms 1/2 (Pkm1/2 transcript, PKM1/2 protein) are key metabolic enzymes that not only link glycolysis and OXPHOS metabolism together but are also hallmark factors in aerobic glycolysis. Pkm1/2 is an allosterically regulated and alternatively spliced gene that produces the pyruvate kinase enzyme responsible for the catalysis of a phosphoryl group from phosphoenolpyruvate in glycolysis to form pyruvate, and the phosphoryl group is transferred to adenosine diphosphate to form ATP (22,23). PKM2 is implicated in cancer and recently PKM1 has been shown to contribute to small cell lung cancer disease onset and progression (24-26). PKM2 reduces OXPHOS when nuclear translocated through phosphorylation, and upregulates glycolytic activity favoring lactate production of acetyl Co-A over a mitochondrial OXPHOS fate – a hallmark of aerobic glycolysis (27,28). Conversely, PKM1 is expressed in tissues requiring large energy requirements, and preferentially promotes pyruvate towards an OXPHOS fate (29). These two isozymes differ as PKM2 has allosteric binding site for fructose-1-6-bisphosphate, when phosphorylated PKM2 dissociated from its highly active homotetrameric conformation into a homodimer for nuclear translocation, whereas PKM1 appears to stably maintain a highly active homotetrameric conformation (24).

As PKM1/2 are implicated in aerobic glycolysis and proliferation, it is important and necessary to investigate their contributions to naïve and primed cell pluripotency. Previous attempts to study the role of PKM1/2 in naïve and primed pluripotent states did not consider the intermediate, formative phases of the pluripotent continuum (30-32). Our study utilized steric-blocking morpholinos to modulate PKM1 and PKM2 isoform levels in naïve and primed mouse pluripotent stem cells and during chemical transitioning of naïve mESCS to formative and primed-like stem cells. The outcomes include effects to Pkm1 and Pkm2 transcript abundance in naïve, formative, primed-like, and primed pluripotent stem cells, and the impact of modulating Pkm1/2 on metabolic and pluripotent state. We also determined the impact of altering Pkm1/2 during transitioning mESCs-to-formative and formative-to-primed-like pluripotency. Our outcomes demonstrate that downregulation of PKM2 alone through splice modifications results in altered metabolic transcript abundance and promotes naïve-to-primed transitioning to formative mEpiLCs. Furthermore, downregulation of PKM2 and upregulation of PKM1 results in a new population of naïve and primed marker expressing cells in primed-like mEpiLCs. This study promotes metabolism as a driver of pluripotency and development, demonstrates how to delineate intermediate states from ground and primed pluripotency and provides evidence of PKM1 having a role in the pluripotent developmental continuum.

## Materials and Methods

### Pluripotent stem cell culture

Mouse embryonic stem cells (mESCs, R1 strain – 129X1 x 129S1 (gifted from Dr. Janet Rossant, The Hospital for Sick Children, Toronto, Ontario, Canada), formative and primed-like mouse epiblast-like cells (mEpiLCs, chemically converted R1 mESCs over 48 and 96 hours), and primed mouse epiblast stem cells (mEpiSCs, strain – 129S2 ((gifted from Dr. Janet Rossant, The Hospital for Sick Children, Toronto, ON), were cultured in the following base media: a 1:1 mixture of KnockOut DMEM/F12 (Thermo Fisher Scientific 12660012) and Neurobasal Media (Thermo Fisher Scientific 21103049) with 0.1 % 2-Mercaptoethanol (Gibco 21985-029), 0.25 % GlutaMAX (Thermo Fisher Scientific 35050061), 1.0 % N2 Supplement (100x) (Thermo Fisher Scientific 17502048), and 2.0 % B27 Supplement (50x) (Thermo Fisher Scientific 17504044) (Supp. Fig. 1a). mESCs and mEpiLCs were cultured on 0.1 % porcine gelatin (Sigma-Aldrich G2500) and mEpiSCs and MEFs were cultured on 10 µg/mL/cm^2^ fibronectin (Roche 11051407001). Base media for the culture of mESCs were supplemented with 1000 units/mL ESGRO Recombinant mouse LIF protein (EMD Millipore ESG1107), and 2i small molecule inhibitors: 1 mM PD0325901 (Reagents Direct 39-C68) and 3 mM CHIR99021 (Reagents Direct 27-H76). Base media for the culture of mEpiLCs and mEpiSCs were supplemented with 20 ng/mL Activin A from mouse (Sigma-Aldrich SRP6057) and 12 ng/mL Fgf-2 from mouse (Sigma-Aldrich SRP3038). mESCs were passaged using StemPro^TM^ Accutase^TM^ (Thermo Fisher Scientific A1110501) and centrifuged at 300 x g for 5 minutes. Primed mouse epiblast stem cells were cultured in the base medium and supplements as mEpiLCs were along with a substratum of irradiated mouse embryonic fibroblasts (MEFs). One hour prior to passaging, growth medium was replaced. Passaging was completed using Gentle Cell Dissociation Buffer (GCDB) (Gibco 13151-014) for 5 minutes at room temperature. Lifted cells were then centrifuged at 244 g for 3 minutes and plated at a seeding density of 1:12 onto MEFs. RNA and protein abundance studies were completed by excluding MEFs for feeder-free conditions and passaging mEpiSCs once with StemPro^TM^ Accutase^TM^ (Thermo Fisher Scientific A1110501) followed by a GCDB passage, this resulted in a clean and healthy population of feeder-free mEpiSCs ready for transcript and protein abundance studies. This study was carried out in biological triplicates.

**FIG 1.**
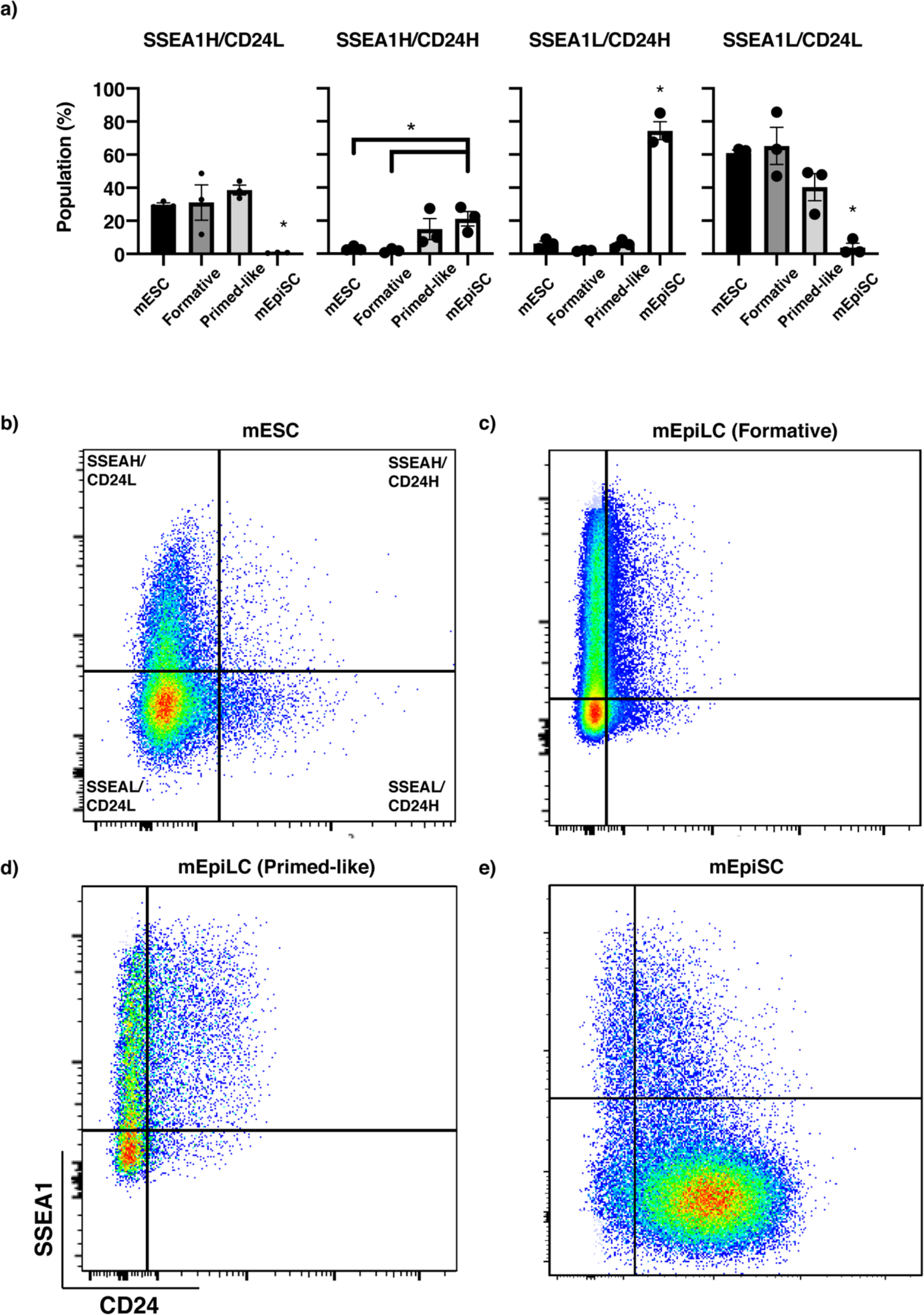
mESC, mEpiLC (Formative), mEpiLC (Primed-like), and mEpiSC SSEA1 and CD24 cell surface marker characterization. Delineation of naïve mESCs, formative and primed-like mEpiLCs, and mEpiSCs by a) flow cytometric analysis of SSEA1 and CD24 cell surface marker expression. Biexponential scale flow plots represent portrayals of SSEA1-compensated Brilliant Violet 421 on a 450 nm laser versus CD24-compensated APC on a 670 nm laser of b) mESCs, c) formative mEpiLC, d) primed-like mEpiLC, and e) mEpiSC. Data shown as mean±SEM of treatments compared in biological triplicate as a one-way ANOVA with a Tukey’s multiple comparisons relative to the control of each corresponding cell type, *p<0.05, n=3 biological replicates.

### Morpholino (MO) delivery into mouse pluripotent stem cells (mPSC)

Splice modifying morpholino oligonucleotides designed to target exon 9 and exon 10 of the PKM gene, where exon 9 is present for Pkm1 and exon 10 is present for Pkm2 (Gene Tools LLC, Philomath, OR) were transfected into mESCs, formative mEpiLCs and mEpiSCs through the scrape delivery method (33). In brief, once cells achieved approximately 70-80% confluency, fresh pluripotent specific media supplemented with 5, 10, or 20 μM morpholino (MO) (fluorescein-tagged control, PKM MO 1 or PKM MO 2 morpholino) that had been 2 μm filter sterilized was ejected onto a PBS(+/+) washed growth surface (either gelatin or fibronectin). The MO supplemented media was swirled for 10 seconds both clockwise and counterclockwise before being allowed to incubate at room temperature for 1 minute. Rubber policeman cell scrapers (Sarstedt 83.3951) were used vertically across the plate, then perpendicularly. Cells co-endocytosed the morpholinos through now open transient pores for 10 minutes without permitting the cells to reattach to their substratum. Transfected cells were replated onto larger growth spaces and allowed to incubate for 24 hours before downstream applications including fluorescent imaging, immunoblotting, transcript abundance and flow cytometry studies. Transfected cells were compared by phase contrast to determine if morphology was influenced. Imaging of fluorescently transfected cells and phase contrast microscopy was completed using a Leica DMI 6000B. Morpholino design, targeting sites and post-transfection changes to PKM1/2 protein can be found in Table 1. Experimental timelines for morpholino transfection and cell fate transitioning are detailed in Supp. Fig. 1.a-b.

**Table 1.**
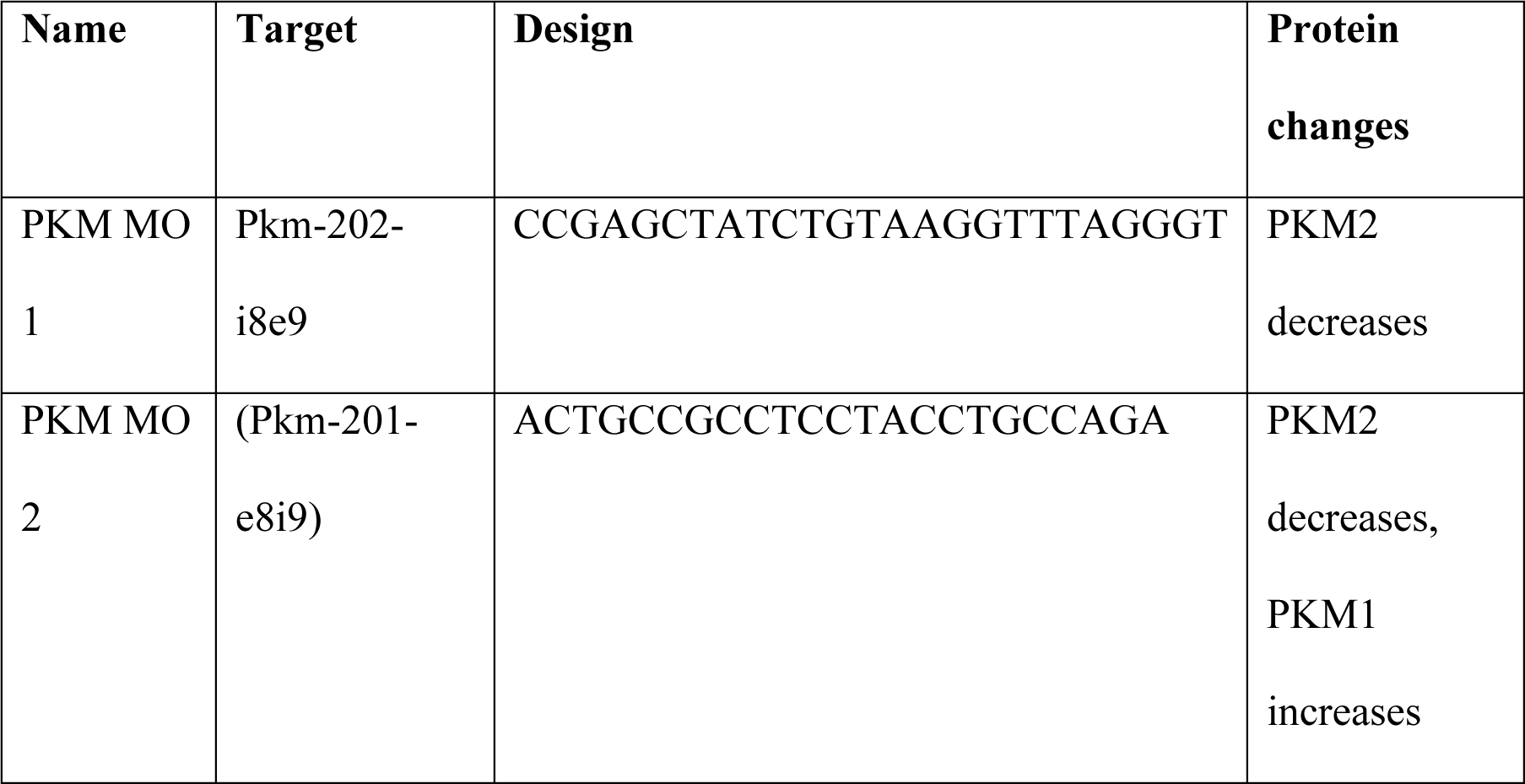
Morpholino Design.

### Quantification of transcript abundance by RT-qPCR

RNA isolation was completed using a RNeasy RNA isolation kit (Qiagen 74104) and Trizol (Ambion 15596018) hybrid protocol followed by DNAse treatment (Invitrogen AM1906). cDNA synthesis was completed using iScript (BioRad 170-8891) on 500 ng of total RNA. Quantitative RT-qPCR was completed using SensiFAST™ SYBR® No-ROX Kit (FroggaBio BIO-98020). Optimal annealing temperatures for each primer were tested in temperature gradients followed by a dilution series to determine primer efficiencies. Relative transcript abundance was calculated using the Pfaffl method of quantification, normalized to mESCs not treated with a morpholino and relative to *α*-Tubulin transcript abundance (34). Forward and reverse primer designs and annealing temperatures are available in Table 2. TaqMan PCR was completed using TaqMan™ Advanced Master Mix (Thermo Fisher Scientific 4444557).

**Table 2.**
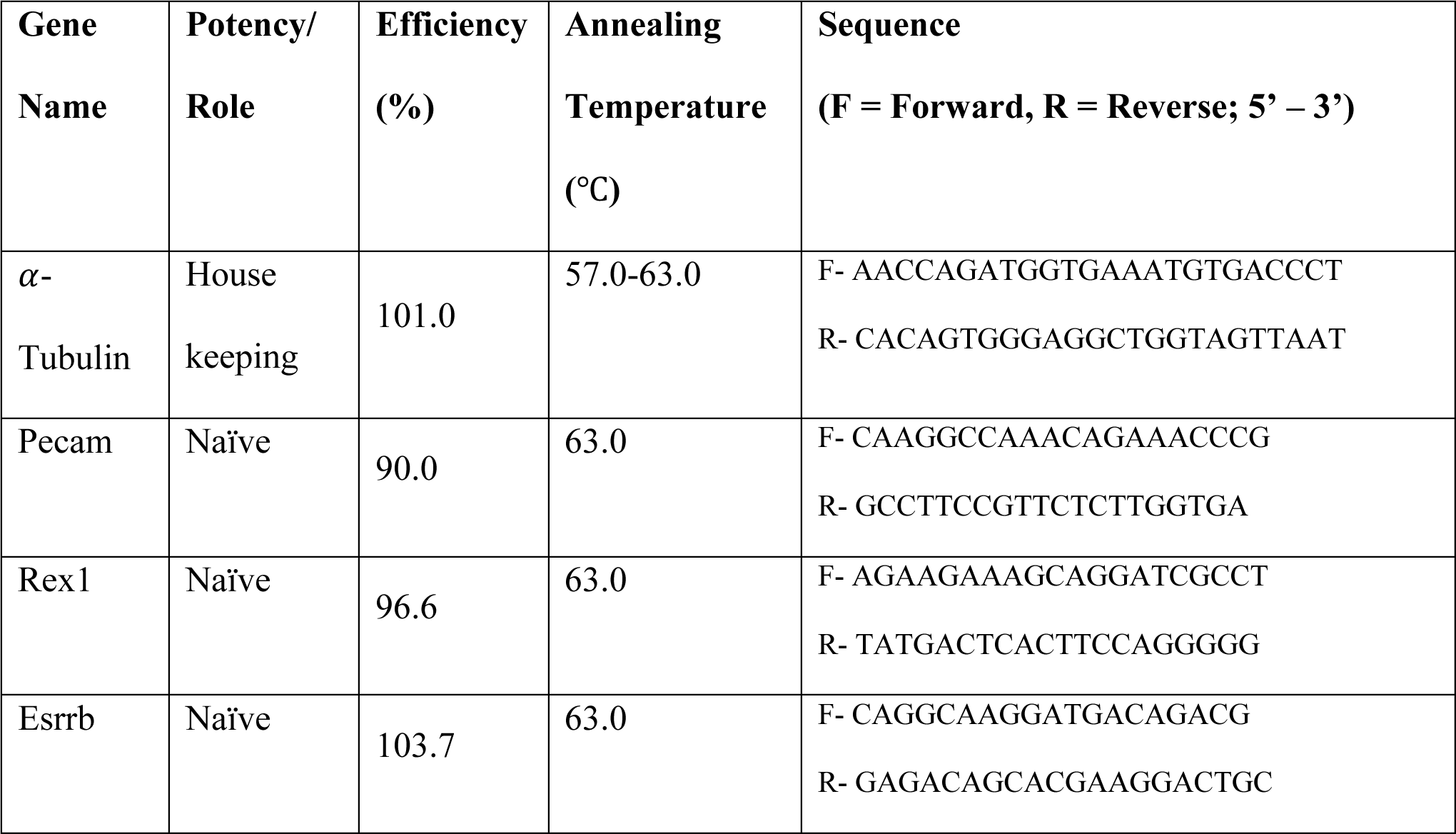
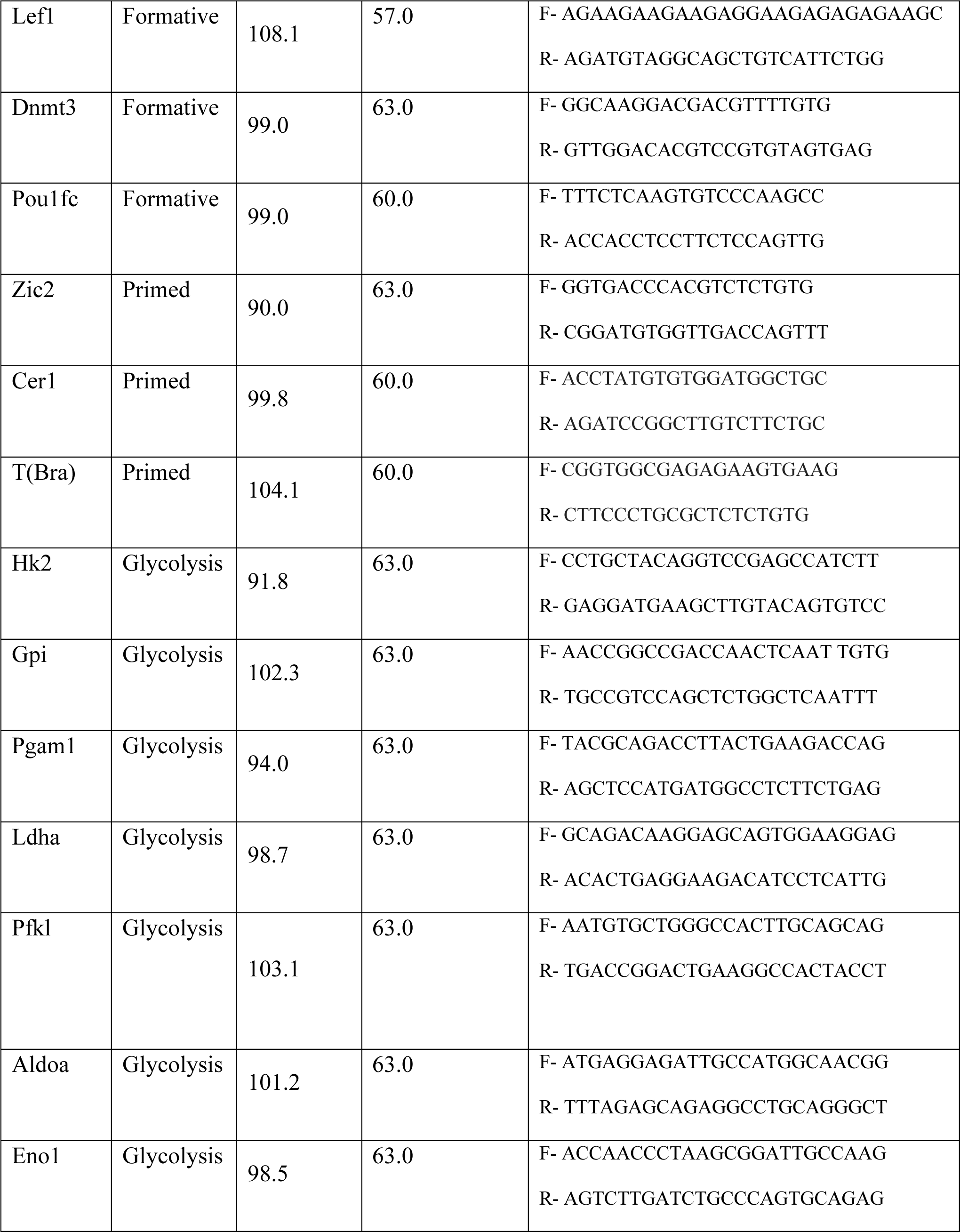
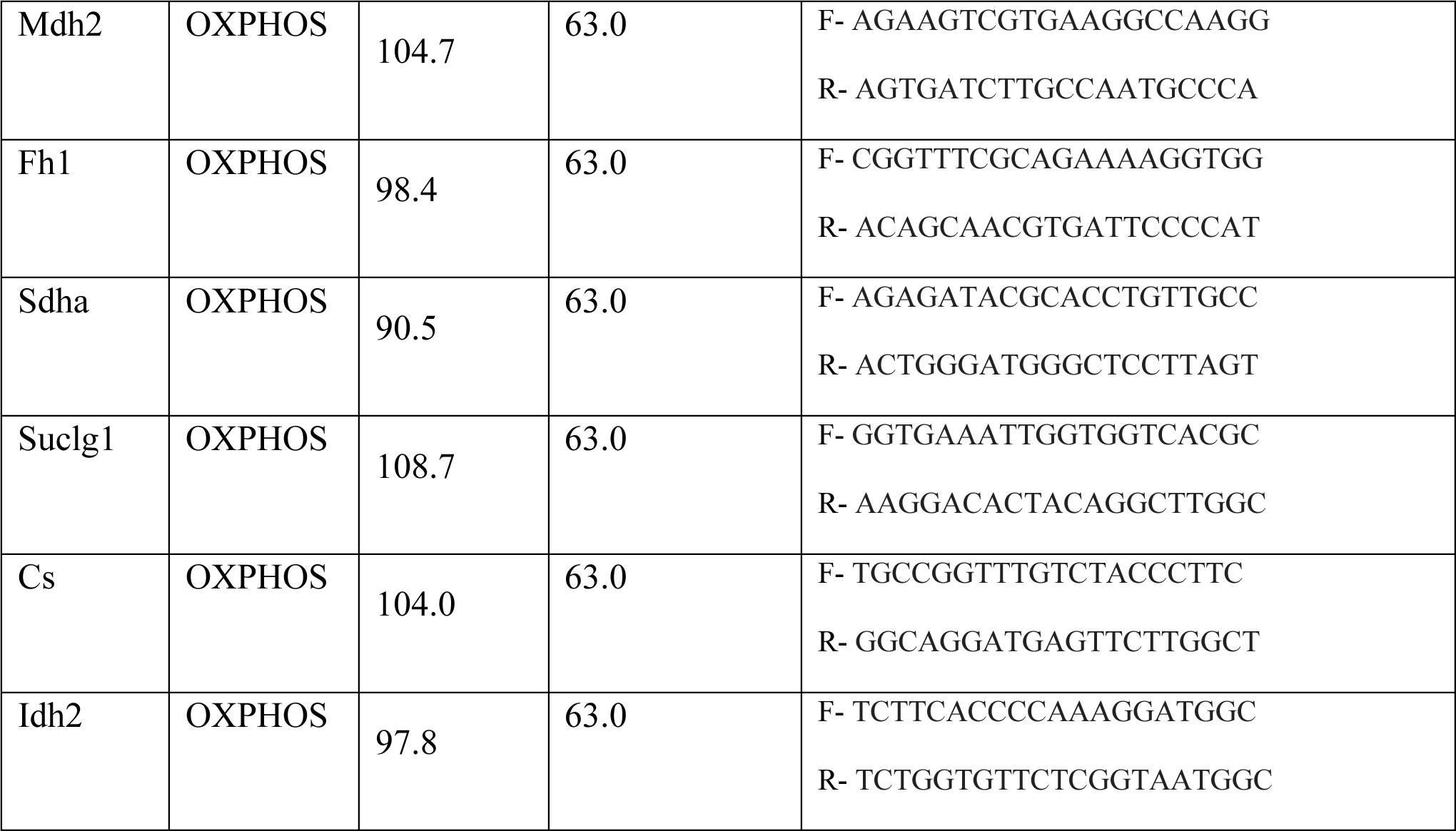
PCR Primers.

Relative transcript abundance was calculated using the ΔΔCt method of quantification, normalized to mESCs not treated with a morpholino and relative to Hprt transcript abundance (35). The fold change in transcript abundance levels for both qPCR and TaqMan studies was calculated and expressed as log2 completed in biological triplicate. MO treated mPSC transcript abundance is relative to their untreated mESCs, mEpiLCs, and mEpiSCs counterpart.

### Quantification of protein levels by Western blot analysis

Cells were washed with chilled (PBS(+/+)) (Gibco 14040-133) and lysed with Pierce^TM^ RIPA Buffer (Thermo Fisher Scientific 89900) supplemented with 1x Phosphatase Inhibitor Cocktail Set 2 (Calbiochem 5246251) and 1x Protease Inhibitor Cocktail Set 1 (Calbiochem 539131).

Protein quantification was completed using a Pierce^TM^ BCA Protein Assay kit (Thermo Fisher Scientific 23225). Loading mixes were prepared at 20 µg in MilliQ H2O, LDS (NuPAGE™ LDS Sample Buffer (4X) Invitrogen NP0007) and Reducing Agent (NuPAGE™ Sample Reducing Agent (10X) Invitrogen NP0004) at 70°C for 10 minutes before loading in NuPAGE^TM^ 4-12% Bis-Tris Gels (Invitrogen NuPAGE NP0336). 1x MOPS (BOLT Invitrogen B000102) and 500 µL of antioxidant containing dithiothreitol (Thermo Fisher Scientific NP0009) was added and electrophoresis was completed at 200V for 50 minutes. Proteins were transferred to a PVDF membrane at 100V over 2 hours. The protein transferred PVDF membrane (EMD Immobilon IPVH00010) was blocked in 5 % bovine serum albumin (BSA) (ALB001) for pPKM2 and 5 % skimmed milk (Carnation) for PKM1 and PKM2 in 1x Tris-Buffered Saline with 0.1 % Tween20 for 1 hour at room temperature with end-to-end agitation. Primary antibodies were incubated overnight at 4 °C with end-to-end rotation. HRP-conjugated secondary antibodies were incubated for 1 hour at room temperature with end-to-end rotation. Membranes were imaged with Luminata Classico Western HRP Substrate (EMD WBLUC0500) and stripped using Restore Western Blot Stripping Buffer (Thermo Fisher Scientific 21059). Bands of interest were compared to *β*-ACTIN. Primary and secondary antibodies and their concentrations are listed in Table 3. Statistics were run in biological triplicate relative to mESCs untreated with a control MO.

**Table 3.** Western blot antibody/marker list.

| **Primary**<br>**Antibody/Marker** | **Concentration** | **Secondary**<br>**Antibody** | **Concentration** |
| --- | --- | --- | --- |
| β-ACTIN<br>A3854 | 1:50000 | N/A: HRP-linked | N/A: HRP-linked |
| PKM1<br>15821-1-AP | 1:5000,<br>5 % TBST | Anti-rabbit IgG,<br>HRP-linked<br>Antibody #7074<br>Cell Signaling<br>Technology | 1:2000-10,000,<br>5 % Milk in<br>TBST |
| pPKM2<br>(Tyr105)<br>3827S | 1:5000,<br>5 % BSA in<br>TBST | Anti-rabbit IgG,<br>HRP-linked<br>Antibody #7074<br>Cell Signaling<br>Technology | 1:2000-10,000,<br>5 % BSA in<br>TBST |
| PKM2<br>15822-1-AP | 1:5000,<br>5 % Skim milk<br>in TBST | Anti-rabbit IgG,<br>HRP-linked<br>Antibody #7074<br>Cell Signaling<br>Technology | 1:2000<br>5 % Milk in<br>TBST |

### Quantification of PSC populations by flow cytometry

Each mPSCs population (mESC, mEpiLC, mEpiSC) were lifted with StemPro^TM^ Accutase^TM^ (Thermo Fisher Scientific A1110501) incubated at 37 °C for 5 minutes. Centrifugation steps were completed at 244 g for 3 minutes. Dead cell compensation and gating was completed using a 50 % mixture of live and dead mESCs stained with Zombie Aqua™ Fixable Viability Kit (BioLegend 423101) and incubated in the light-tight container at room temperature for 30 minutes. Cells were washed with 2 mL of flow cytometry staining buffer (FCSB) containing: 90 % PBS(-/-), 10 % FBS (qualified, ESC grade), and fixed with 4 % paraformaldyde (PFA) in PBS(-/-). Fixed cells were washed with PBS(-/-), centrifuged and divided into unstained, single, full-minus-one and full stained combinations of each cell type. Fixed cells were stained with conjugated antibodies for 1 hour in a light-tight box at room temperature prior to washing, centrifugation and resuspension in PBS(-/-) and ejected through a 40 µm cell strainer (Fisherbrand™ Sterile Cell Strainers 22-363-547) with a final wash of 100 µL of PBS(-/-). Flow cytometry was completed on a FACSCanto flow cytometer. Antibodies and their concentrations are listed in Table 4. Treatments and controls were run in biological triplicate, MO treated mPSC cell surface marker events are relative to their corresponding mESC, mEpiLC, and mEpiSC counterpart.

**Table 4.** Flow cytometry antibody/marker list.

| **Primary**<br>**Antibody/Marker** | **Concentration** | **Secondary**<br>**Antibody** | **Concentration** |
| --- | --- | --- | --- |
| CD24<br>138505 | 0.25 µg/1<br>million cells | N/A:<br>Conjugated | N/A:<br>Conjugated |
| SSEA1<br>125614 | 5 µL/1 million<br>cells | N/A:<br>Conjugated | N/A:<br>Conjugated |

### Statistical Analyses

Statistics were completed using a one- and two-way ANOVAs where applicable. Characterization by flow cytometry of SSEA1/CD24 in mPSCs was completed using a one-way ANOVA with Tukey’s multiple comparisons test. Flow cytometric analysis of transfection efficiency compared random control morpholino groups using a one-way ANOVA with Tukey’s multiple comparisons test. Immunoblotting for protein abundance in the influence of PKM1/2 morpholinos utilized a one-way ANOVA with a Dunnett’s multiple comparisons test.

Determining the influence of PKM1/2 morpholinos on transitioning formative and primed-like mEpiLCs as quantified by flow cytometry of SSEA1/CD24 in mPSCs was completed using a one-way ANOVA with Dunnett’s multiple comparisons test. Transcript abundance studies examining the influence of PKM1/2 morpholinos on mPSCs was accomplished using a two-way ANOVA with Dunnett’s multiple comparisons test with significance at P<0.05.

## Results

### Formative and primed-like mEpiLCs can be distinguished from primed mEpiSCs through SSEA1 and CD24 cell surface expression

mEpiLCs can be distinguished from naïve mESCs and primed mEpiSCs based on Stage - Specific Embryonic Antigen-1 (SSEA1) and Cluster of differentiation 24 (CD24) expression (Fig. 1.a-e). Representative flow cytometry plots demonstrate cell population quantification (%) of pluripotent cells with high and low expression of SSEA1 and CD24 (Fig. 1.b-e). There was a significant difference between group mean values of pluripotent cell types expressing high levels of SSEA1 and low levels of CD24 (F(3,8)=8.993, p=0.0061) (Fig. 1.a). Tukey’s multiple comparisons post hoc test determined that the percentage of mEpiSCs expressing high levels of SSEA1 and low levels of CD24 was significantly greater than mESCs, mEpiLCs (formative), or mEpiLCs (primed-like) (p=0.0253, p=0200, p=0.0058 respectively) (Fig. 1.a). Moreover, there was a significant difference between group mean values of pluripotent cell types expressing high levels of SSEA1 and CD24 (F(3,8)=5.777, p=0.0212) (Fig. 1.a). Tukey’s multiple comparisons post hoc test determined that the percentage of mEpiSCs expressing high levels of SSEA1 and CD24 was significantly greater than mESCs or mEpiLCs (formative) (p=0.0458, p=0.0319 respectively) (Fig. 1.a). There was also a significant difference between group mean values of pluripotent cell types expressing low levels of SSEA1 and high levels of CD24 (F(3,8)=142.9, p<0.0001) (Fig. 1.a). Tukey’s multiple comparisons test determined that the percentage of mEpiSCs expressing low levels of SSEA1 and high levels of CD24 was significantly greater than mESCs, mEpiLCs (formative), or mEpiLCs (primed-like) (p=<0.0001 in all instances) (Fig. 1.a). Lastly, there was a significant difference between group mean values of pluripotent cell types expressing low levels of SSEA1 and CD24 (F(3,8)=15.54, p<0.0011) (Fig. 1.a). Tukey’s multiple comparisons test determined that the percentage of mEpiSCs expressing low levels of SSEA1 and CD24 was significantly greater than mESCs, mEpiLCs (formative), or mEpiLCs (primed-like) (p=0.0021, p=0013, p=0.0276 respectively) (Fig. 1.a).

### Efficient transfection of morpholino oligonucleotides into mESCs

Random control Morpholinos tagged with a fluorescein label were scrape delivered into mESCs as a concentration series of 5, 10, or 20 μM (Fig. 2.a-c). At 10 μM, tagged control morpholinos were detectable by fluorescent microscopy (Fig. 2.a). Scrape delivered cells were measured via flow cytometry for transfection efficiency as assessed by FITC+ events relative to the total live cell population and mean fluorescence intensity as measured by geometric mean of fluorescent events (Fig. 2.b, c). There was a significant concentration-dependent difference between FITC+ cell events using random control morpholino treatments as determined by a one-way ANOVA (F (2,6)=45.77, p=0.0002) (Fig. 2.b). The concentration series for FITC+ events demonstrated: 5 μM (mean=73.9%, SEM=2.1), 10 μM (mean=94.7%, SEM=2.3), and 20 μM (mean=98.0, SEM=1.3) (Fig. 2.b).

**FIG 2.**
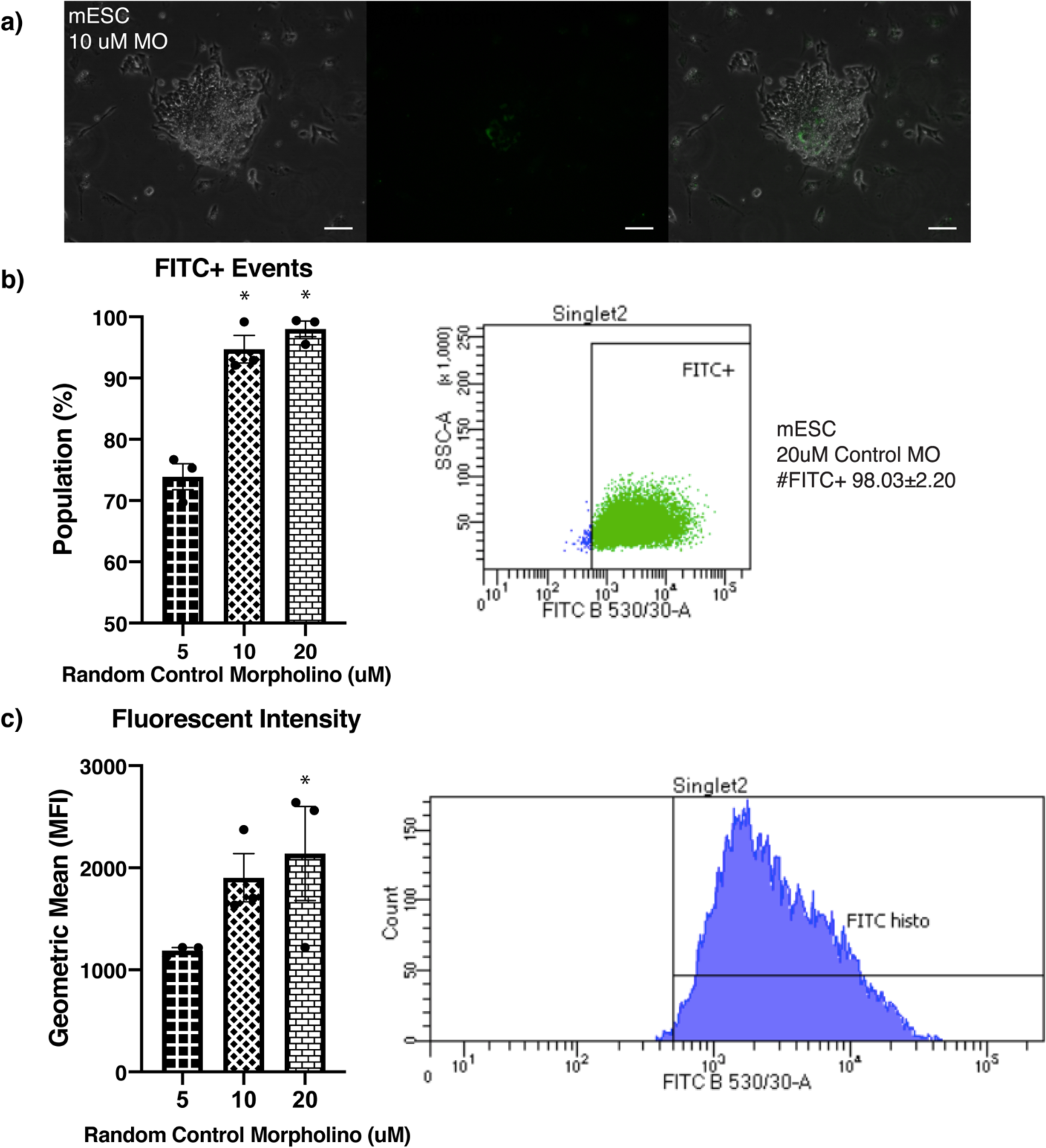
mESCs can be efficiently and effectively transfected with morpholinos by scrape delivery at 20 μM. Scrape delivered morpholinos tagged with a fluorescein label were visible in a) mESCs as demonstrated by fluorescent phase contrast microscopy at concentration of 10 μM. Scale bars represent 75 μm. Transfection of morpholinos was optimized by comparing 5, 10, and b) 20 μM fluorescein-tagged control morpholinos as determined through c) FITC+ frequency relative to total population and d) geometric mean by flow cytometric analysis of FITC wavelength laser channel. Data shown as mean±SEM of treatments compared in biological triplicate, *p<0.05, n=3 biological replicates.

### Pkm1/2 transcript abundance is altered following PKM1/2 spliceosome modification

A two-way ANOVA was conducted to examine the influence of treating each pluripotent state with the PKM MO 1 or PKM MO 2 morpholinos at 20 μM on Pkm1/2 transcript abundance levels (Fig. 3.a-c). There was a statistically significant interaction, suggesting the effects on different pluripotent cell types varies with the morpholino treatment, for Pkm1 (Fig. 3.a) and Pkm2 (Fig. 3.b) transcript abundance, but not the Pkm1/Pkm2 (Fig. 3.c) transcript abundance ratio, (F(6,24)=11.92, p<0.001), (F(6,24)=2.695, p=0.0382), and (F(6,24)=1.904, p=0.1214) respectively between pluripotent cell type and morpholino treatment. Simple main effect analysis demonstrated that within pluripotent cell types, the addition of PKM MO 1 or PKM MO 2 morpholinos significantly influenced Pkm1 transcript abundance (F(3,24)=6.253, p=0.0027, and F(2,24)=25.86, p<0.001 respectively). Simple main effect analysis also demonstrated that pluripotent cell type and the addition of PKM MO 1 or PKM MO 2 morpholinos significantly influenced Pkm1/Pkm2 transcript abundance ratio (F(3,24)=14.26, p<0.0001, and F(2,24)=21.79, p<0.0001 respectively). Based on Dunnett’s multiple comparisons tests relative to the control treatment of each pluripotent cell type, adding a PKM MO 2 morpholino significantly enhanced Pkm1 transcript abundance in mESCs (p<0.0001), mEpiLC (formative) (p=0.0006), and mEpiSCs (p=0.0080), and Pkm1/Pkm2 transcript abundance ratio in mESCs (p=0.0023) and mEpiLCs (formative) (p=0.0025). Based on Dunnett’s multiple comparisons tests relative to the control treatment of each pluripotent cell type, treatment with the PKM MO 2 morpholino significantly reduced Pkm1 and Pkm2 transcript abundance in mEpiLC (primed-like) (p=0.0025 and p=0.0497, respectively).

**FIG 3.**
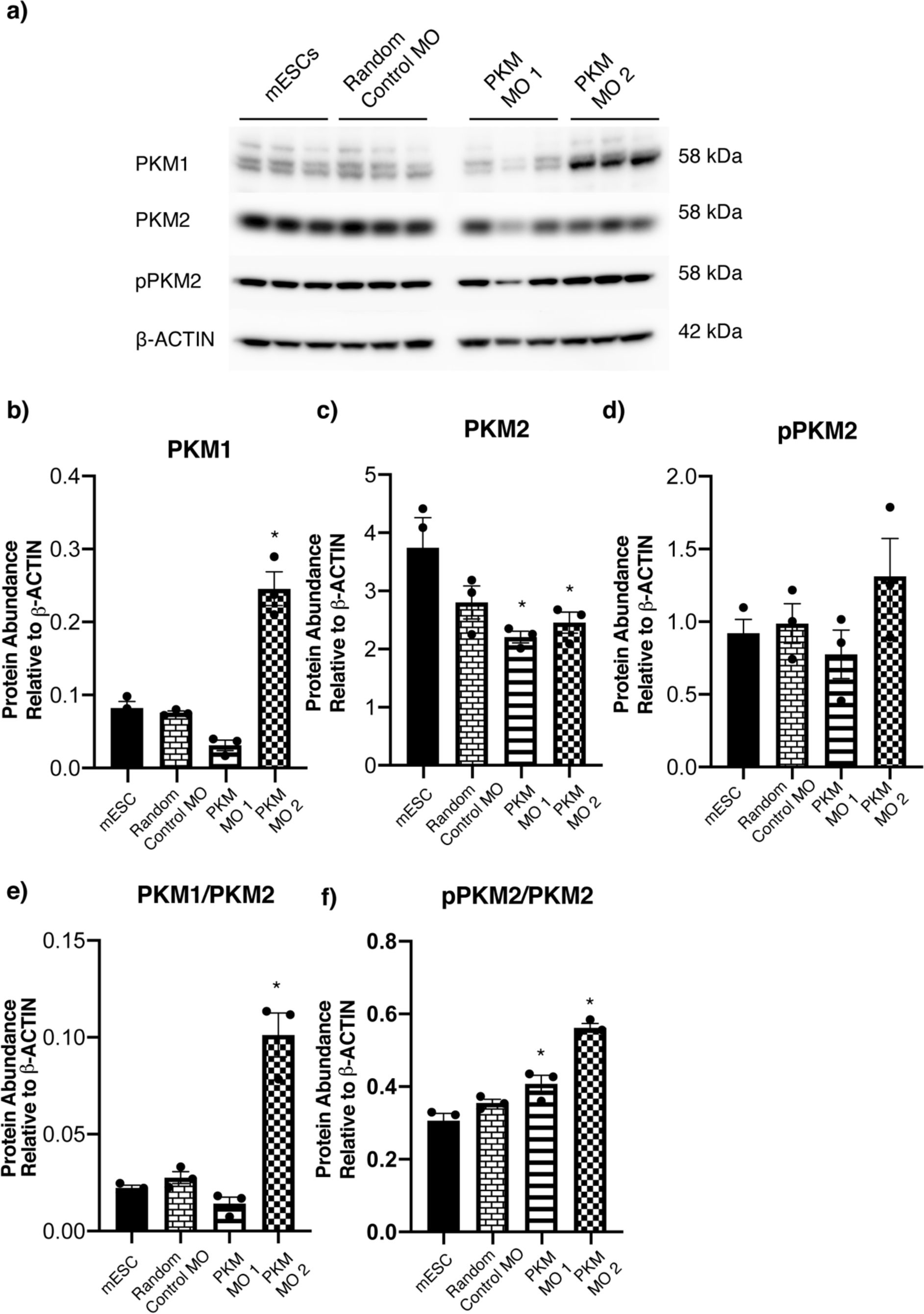
PKM morpholinos influence Pkm1/2 transcript abundance in naïve mESCs, formative mEpiLCs, primed-like mEpiLC, and primed mEpiSCs. Delivery of the PKM MO 2 significantly influences a) Pkm1 transcript abundance in mESCs, formative mEpiLCs, primed-like mEpiLCs, and mEpiSCs relative to control cells of each pluripotent state. b) Pkm2 transcript abundance was significantly downregulated in mEpiLCs with PKM2 morpholino delivery. mESCs and formative mEpiLCs c) Pkm1 to Pkm2 transcript abundance ratio was upregulated following PKM2 morpholino transfection. Data shown as mean±SEM of treatments compared in biological triplicate as a two-way ANOVA with a Dunnett’s multiple comparisons relative to the control of each corresponding cell type, *p<0.05, n=3 biological replicates run in technical triplicate. Data represents Log2 of fold change relative to Hprt and normalized to control mESCs.

### Steric blocking morpholinos alter PKM1 and PKM2 protein levels

Forty-eight hours-post 20 μM morpholino scrape delivery produced significant changes to PKM1 and PKM2 protein levels in mESCs (Fig. 4.a, Supp. Fig. 1.b). The transfection of a random control morpholino did not affect PKM1 or PKM2 protein abundance compared to mESCs scraped without a morpholino (Fig. 4.a-f). There was a significant difference between group mean values of PKM1 protein abundance following morpholino treatments as determined by a one-way ANOVA (F(3,8)=52.21, p<0001) (Fig. 4.b). There was a significant difference between group mean values of PKM2 protein abundance following morpholino treatments as determined by a one-way ANOVA (F (3,8)=4.619, p=0.0371) (Fig. 5.c). A Dunnett’s multiple comparison test determined that the addition of the PKM1 designed Morpholino, now referred to as ‘PKM MO 1’ significantly decreased PKM2 protein abundance (p=0.0212) (Fig. 4.b, c). The PKM2 designed morpholino, now referred to as ‘PKM MO 2’ significantly decreased PKM2 protein abundance (p=0.0480) and significantly increased PKM1 protein abundance (P<0.0001) (Fig. 4.b, c). There was a significant increase in the ratio of PKM1:PKM2 protein abundance with the addition of the PKM MO 2 (p<0.0001) (Fig. 4.e). There was a significant increase in the ratio of PKM1:PKM2 protein abundance with the addition of the PKM MO 1 and the PKM MO 2 (p=0.0082 and p<0.0001) (Fig. 4.e). There was a significant increase in the ratio of pPKM2:PKM2 protein abundance with the addition of the PKM MO 1 and the PKM MO 2 (p=0.0082 and p<0.0001) (Fig. 4.f).

**FIG 4.**
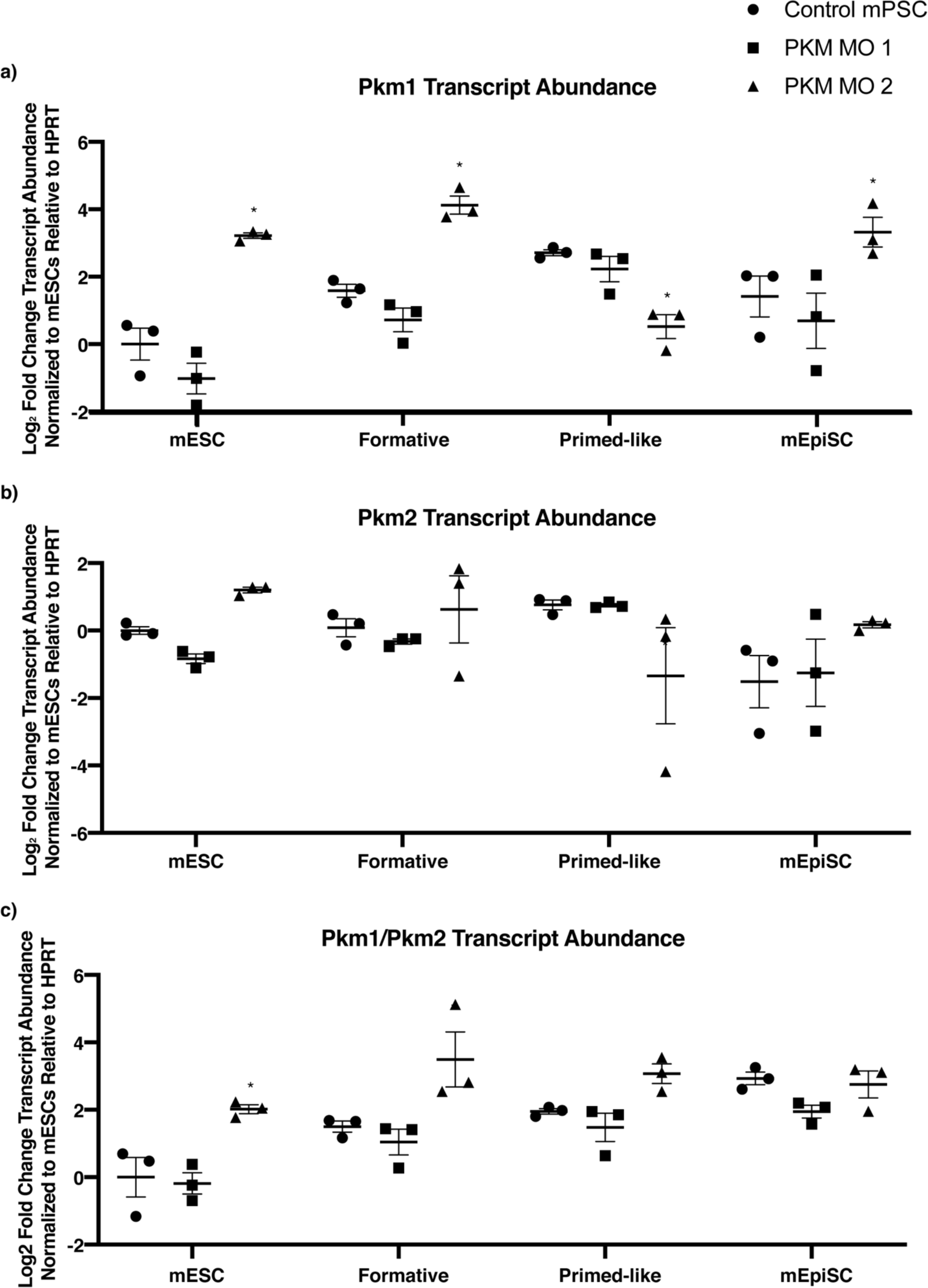
Splice-modifying morpholinos modulate PKM1 and PKM2 protein abundance in mESCs. a) Representative PKM1/2 immunoblotting of morpholino transfected mESCs demonstrates b) upregulation of PKM1 and c) downregulation of PKM2 by transfecting the PKM MO2, whereas PKM MO1 significantly downregulates PKM2. The PKM MO 2 additionally e) upregulates protein abundance of PKM1/PKM2 ratio. There was d) no significant change in pPKM2 with the inclusion of a PKM morpholino. Data shown as mean±SEM of treatments compared in biological triplicate as a one-way ANOVA with a Tukey’s multiple comparisons test, *p<0.05, n=3 biological replicates.

**FIG 5.**
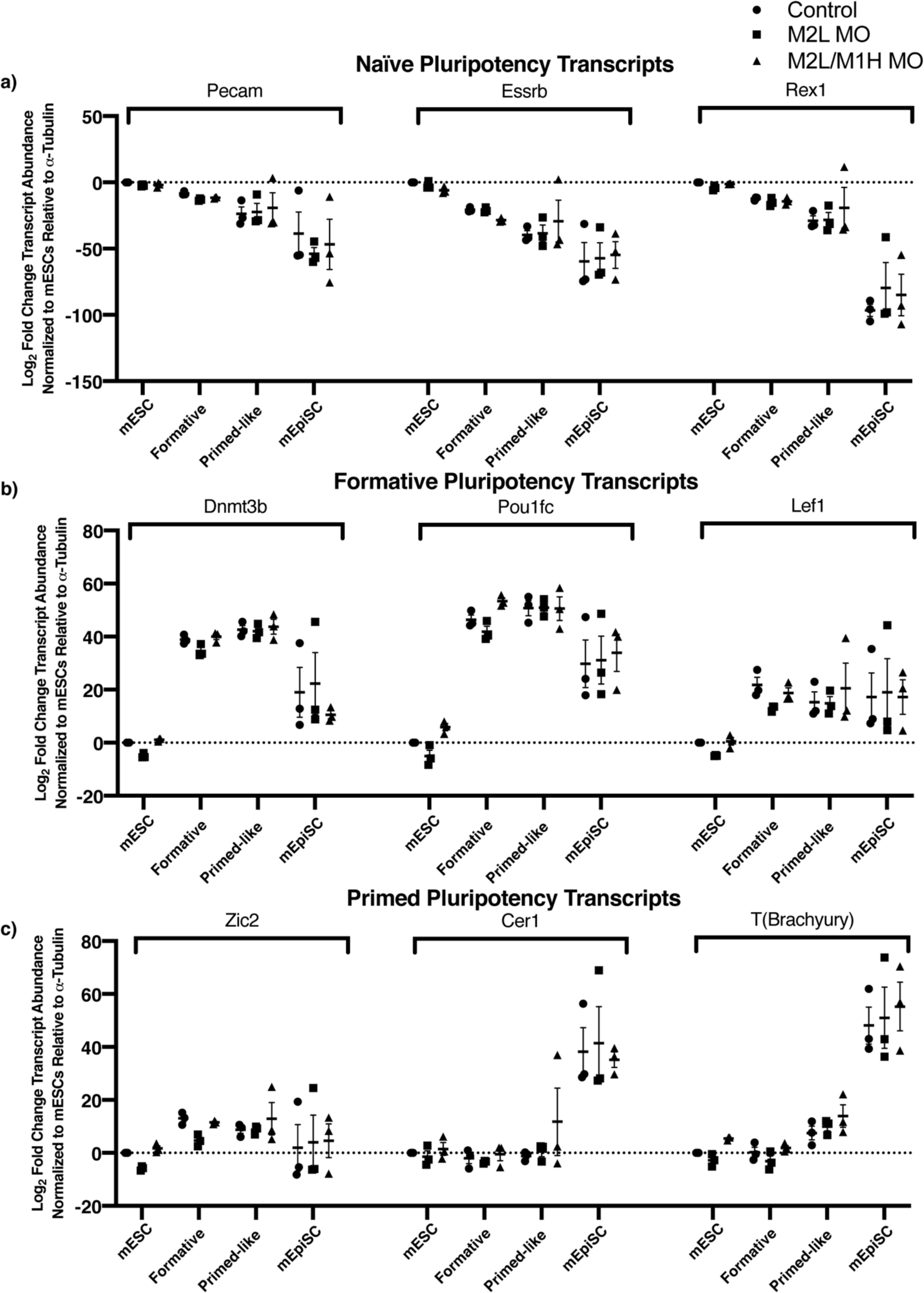
PKM morpholinos do not influence key naïve, formative, and primed pluripotency transcript abundance markers in naïve mESCs, formative mEpiLCs, primed-like mEpiLC, and primed mEpiSCs. Quantification of transcript abundance following scrape delivery of 20μM PKM MO 1 and PKM MO 2 morpholinos into mPSCs over 48 hours. Transcript markers include a) naïve pluripotency associated genes Pecam, Esrrb, and Rex1, b) formative pluripotency associated genes Dnmt3b, Pou1fc, and Lef1, and c) primed pluripotency associated genes Zic2, Cer1, and T(Brachyury). Transcript abundance was compared using the Pfaffl method and data is shown as mean±SEM of treatments compared in biological triplicate as a two-way ANOVA with a Dunnett’s multiple comparisons relative to the control of each corresponding cell type, *p<0.05, n=3 biological replicates run in technical triplicate. Data represents Log2 of fold change relative to *α*-Tubulin and normalized to control mESCs.

### Morpholino-induced decreased PKM2 and increased PKM1 protein abundance decreases transcript abundance of glycolytic genes Eno1 and Hk2 in primed mEpiSCs

A two-way ANOVA was conducted that examined the influence of treatment with PKM MO 1 or PKM MO 2 morpholinos (20 μM) on glycolytic and oxidative phosphorylation (OXPHOS) metabolic transcript abundance following transfection in mESCs, transitioning mEpiLCs to the formative and primed-like pluripotent states and mEpiSCs over 48 hours. There was no significant interaction for the glycolysis genes Hexokinase 2 (Hk2), Lactate dehydrogenase A (Ldha), Phosphofructokinase 1 (Pfk1), and Alpha-enolase (Eno1) transcript abundance (Fig. 6.a), however, there was a statistically significant difference in transcript abundance between pluripotent cell types (F3,24)=38.12,p<0.0001), (F(3,24)=13.80, p<0.0001), (F(3,24)=5.361, p=0.0057), and (F(3,24)=4.815, p<0.0092) respectively. Based on Dunnett’s multiple comparisons tests relative to the control treatment of each pluripotent cell type, treatment with the PKM MO 2 morpholino significantly reduced Eno1 and Hk2 transcript abundance in mEpiSCs (p=0.0268 and p=0.0128 respectively). There was no significant interaction for the OXPHOS genes Isocitrate dehydrogenase 2 (Idh2), malate dehydrogenase 2 (Mdh2), and Succinate-CoA ligase (Suclg1) transcript abundance (Fig. 6.b), however, there was a statistically significant difference in transcript abundance between pluripotent cell types (F(3,24)=10.35, p=0.0001),(F(3,24)=6.679, p=0.0019), and (F(3,24)=3.299, p=0.0375).

**FIG 6.**
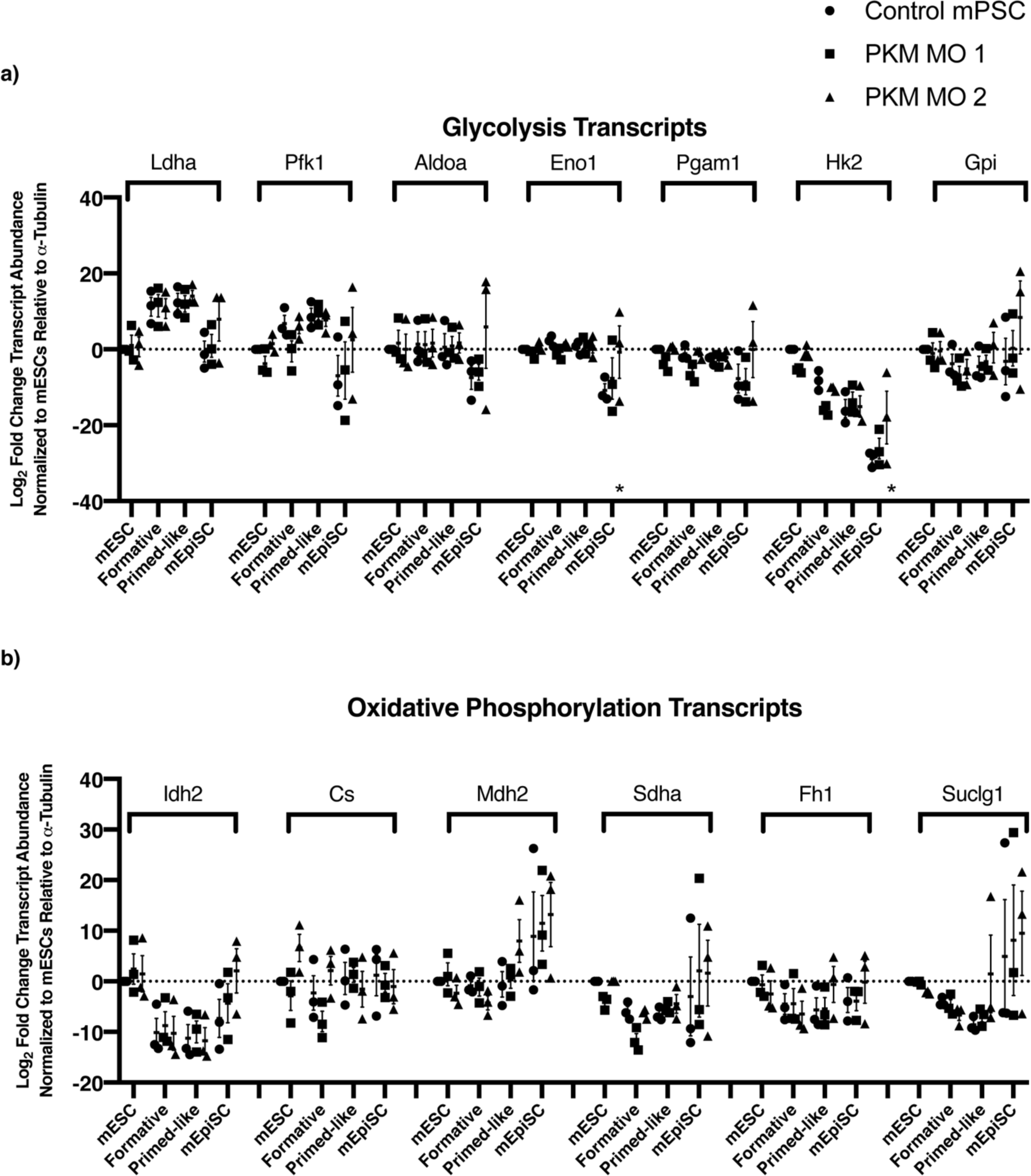
PKM morpholinos influence key glycolytic transcript abundance markers in naïve mESCs, formative mEpiLCs, primed-like mEpiLC, and primed mEpiSCs. Quantification of transcript abundance following scrape delivery of 20 μM PKM MO 1 and PKM MO 2 morpholinos into mPSCs over 48 hours. Transcript markers include a) glycolysis genes Hk2, Gpi, Pfkl, Aldoa, Pgam1, Eno1, and Ldha, and b) oxidative phosphorylation genes Cs, Idh2, Suclg2, Sdh-a, Fh, and Mdh2. Transcript abundance was compared using the Pfaffl method and data is shown as mean±SEM of treatments compared in biological triplicate as a two-way ANOVA with a Dunnett’s multiple comparisons relative to the control of each corresponding cell type, *p<0.05, n=3 biological replicates run in technical triplicate. Data represents Log2 of fold change relative to *α*-Tubulin and normalized to control mESCs.

### PKM1/2 modulation does not alter naïve, formative, or primed pluripotency associated transcripts

A two-way ANOVA was conducted that examined the influence of treating with the PKM MO 1 or PKM MO 2 morpholinos on transcript abundance of naïve, formative, and primed pluripotent associated transcripts. There was no significant interaction for the naïve pluripotency genes Rex1, Pecam, or Esrrb transcript abundance (Fig. 6.a), however, there was a statistically significant difference in transcript abundance between pluripotent cell types (F(3,24)=54.98, p<0.0001), (F(3,24)=15.93, p<0.0001), and (F(3,24)=25.06, p<0.0001) respectively. No significant interaction for the formative pluripotency genes Lef1 (Lymphoid Enhancer Binding Factor 1), Dnmt3b, or Pou1fc transcript abundances were observed (Fig. 6.b), however, there was a statistically significant difference in transcript abundance between pluripotent cell types (F(3,24)=7.380, p=0.0011), (F(3,24)=60.28, p<0.0001), and (F(3,24)=85.18, p<0.0001) respectively. No significant interaction for the primed pluripotency genes Zic2, Cer1, or T(Brachyury) transcript abundances was observed (Fig. 6.c), however, a statistically significant difference in transcript abundance was detected between pluripotent cell types (F(3,24)=4.071, p=0.0180), (F(3,24)=27.79, p<0.0001), and (F(3,24)=70.40, p<0.0001) respectively.

### PKM1/2 modification alters SSEA1 and CD24 ratios in transitioning mESCs into formative state and formative mEpiLCs into primed-like state mEpiLCs

We detected a significant difference between group mean values of morpholino treatments expressing low levels of SSEA and high levels of CD24 in formative mEpiLCs as determined by a one-way ANOVA (F (2,6)=8.167, p=0.0194) (Fig. 7.a). A Dunnett’s multiple comparison test detected that the percentage of mEpiLCs (formative) transfected with the PKM MO 1 morpholino (Fig. 7.c) was significantly enhanced compared to control mEpiLCs (p=0.0128).

**FIG 7.**
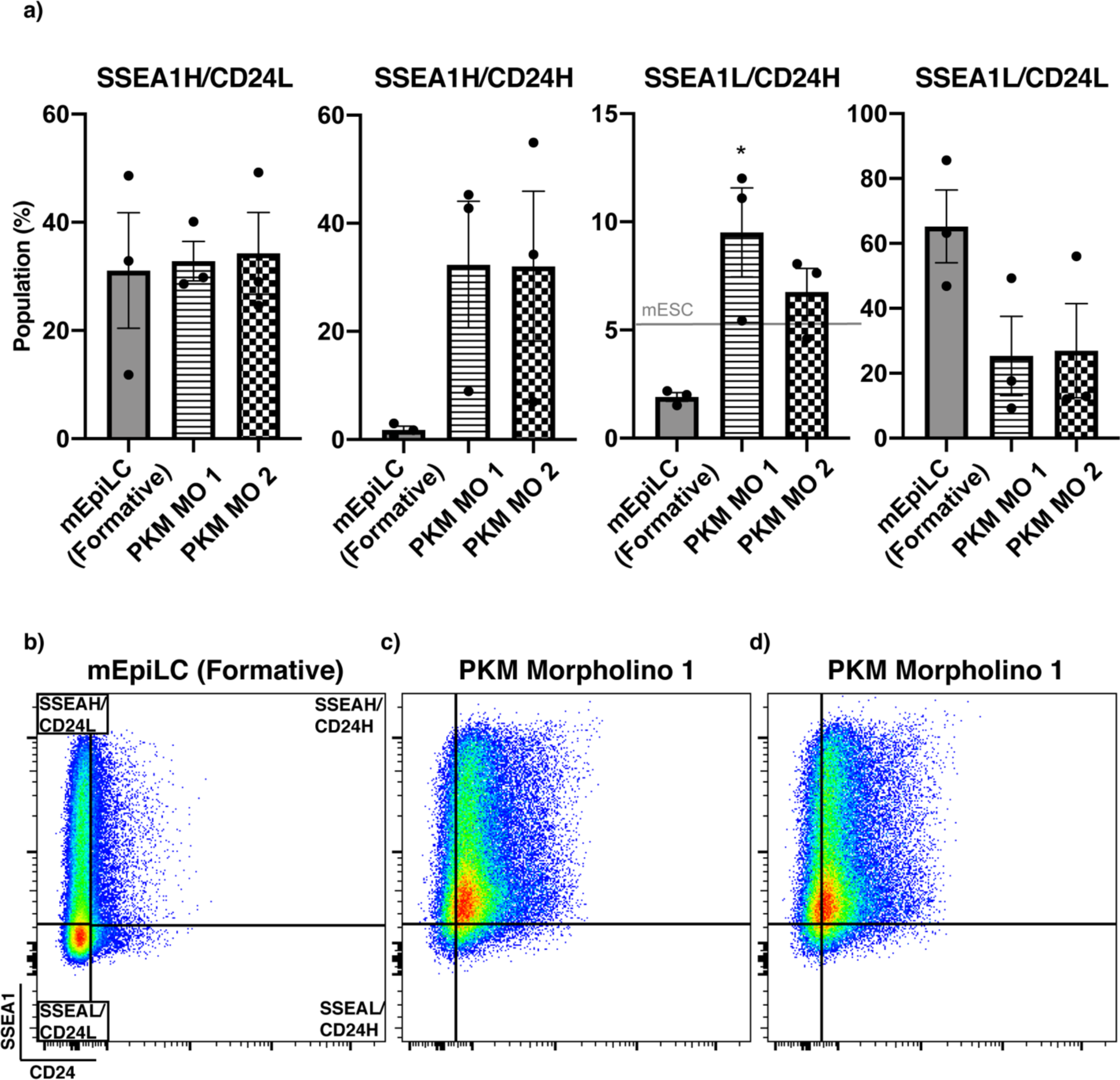
Influence of downregulating PKM on SSEA1 and CD24 expression in transitioning formative mEpiLCs. Scrape delivery of 20 μM PKM MO 1 and PKM MO 2 morpholinos into mESCs transitioned over 48 hours into formative mEpiLCs compared by a) SSEA1 and CD24 cell surface markers by flow cytometry. Transitioning into formative b) mEpiLCs with the addition of c) PKM MO 1 morpholinos and significantly increased the population of SSEA1H and CD24H cells and both PKM MO 1 and d) PKM MO 2 delivery resulted in decreased SSEA1L and CD24L cell populations. Biexponential scale flow plots represent portrayals of SSEA1-compensated Brilliant Violet 421 on a 450 nm laser versus CD24-compensated APC on a 670 nm laser. Data shown as mean±SEM of treatments compared in biological triplicate as a one-way ANOVA with a Dunnett’s multiple comparisons relative to the control of each corresponding cell type, *p<0.05, n=3 biological replicates.

There was an observed significant difference between group mean values of morpholino treatments expressing high levels of SSEA and CD24 in primed-like mEpiLCs as determined by a one-way ANOVA (F(2,6)=8.486, p=0.0178) (Fig. 7.a). A Dunnett’s multiple comparison test determined that the percentage of mEpiLCs (primed-like) transfected with the PKM MO 1 (Fig. 8.c) and PKM MO 2 morpholinos (Fig.8.d) was significantly greater than control mEpiLCs for SSEA1 high and CD24 high events (p=0.0264 and p=0.0172 respectively).

**FIG 8.**
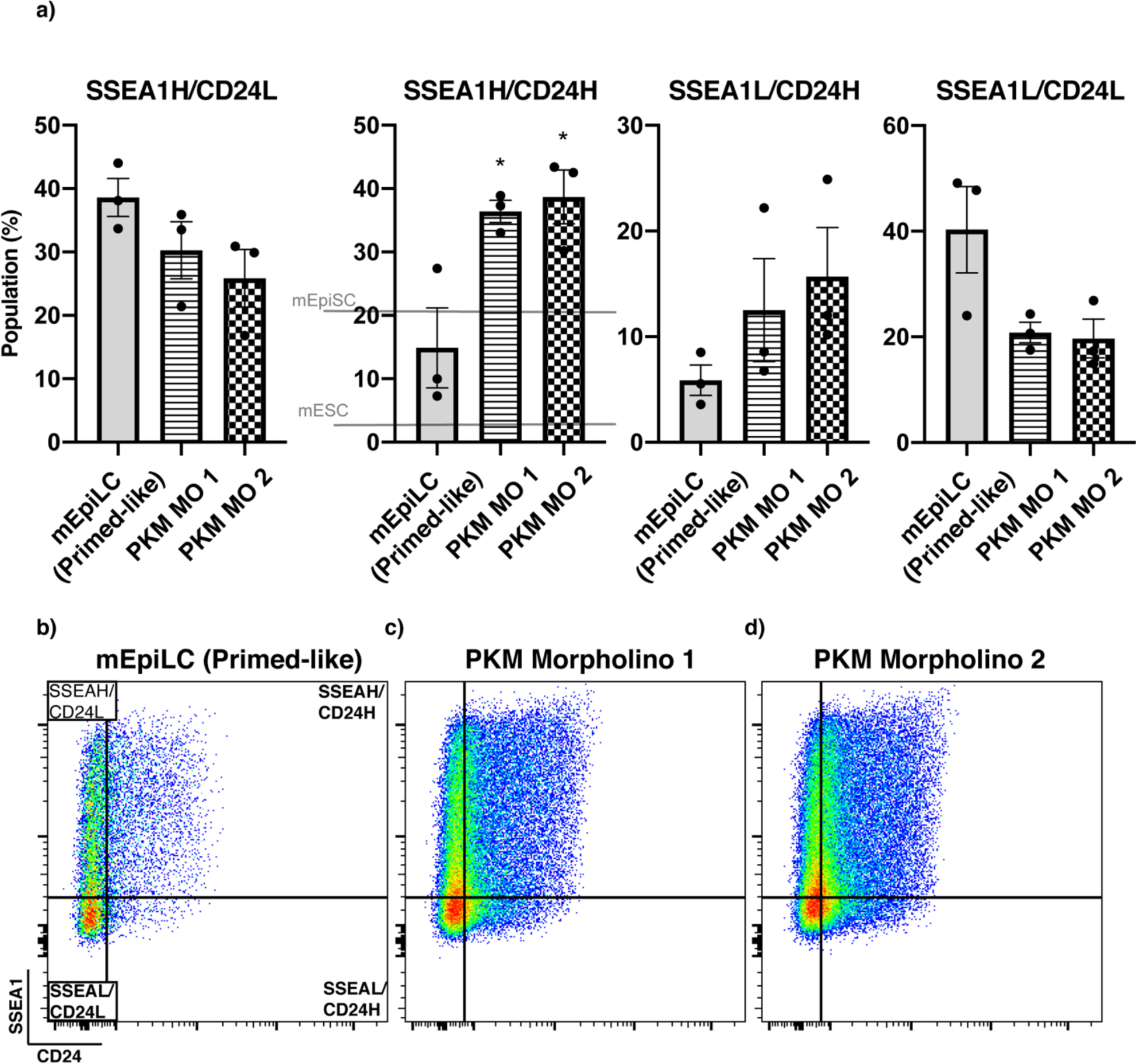
Influence of downregulating PKM on SSEA1 and CD24 expression in transitioning primed-like mEpiLCs. Scrape delivery of 20 μM PKM MO 1 and PKM MO 2 morpholinos into formative mEpiLCs transitioned over 48 hours into primed-like, mEpiLCs compared by a) SSEA1 and CD24 cell surface markers by flow cytometry. Transitioning into primed-like b) mEpiLCs with the addition of c) PKM MO 1 morpholinos and d) PKM MO 2 morpholino delivery significantly increased the population of SSEA1H and CD24H cells and decreased SSEA1L and CD24L cell populations. Biexponential scale flow plots represent portrayals of SSEA1-compensated Brilliant Violet 421 on a 450 nm laser versus CD24-compensated APC on a 670 nm laser. Data shown as mean±SEM of treatments compared in biological triplicate as a one-way ANOVA with a Dunnett’s multiple comparisons relative to the control of each corresponding cell type, *p<0.05, n=3 biological replicates.

## Discussion

Growing evidence is promoting cellular metabolism to having an active role in cell pluripotency, differentiation, and development (16,36,37). Within the cells of the early embryo and the pluripotent continuum, pyruvate kinase muscle isoforms 1 and 2 are suggested to play a variety of key roles (31,38,39). Lentiviral overexpression of either Pkm1 or Pkm2 isoform in mESCs significantly increased the transcript abundance of pluripotency associated genes Nanog, Eras, and Rex1, but did not influence their cell fate when exposed to differentiation media (30). Conversely, when Pkm1/2 were downregulated via shRNAs, Nanog, Eras, and Rex1 were significantly downregulated suggesting PKM1/2 expression maintains the pluripotent state. Indeed, downregulating total Pkm1/2 significantly hinders cellular reprogramming towards pluripotency (30). Overexpression of Pkm2, but not Pkm1, significantly increased alkaline phosphatase staining during reprogramming, suggesting that Pkm2 is the PKM isoform that facilitates iPSC generation. The metabolic switch from OXPHOS to glycolysis is linked to hypoxia inducible factor 1*α*(HIF-1*α*) activation, where PKM2 interacts with the HIF1*α*subunit promoting transactivation domain function as well as p300 recruitment to the HIF1*α*response elements during somatic cell reprogramming (40). This interaction promotes the switch from OXPHOS to aerobic glycolysis (41). The reprogramming of OXPHOS-reliant human somatic cells to iPSCs results in primed pluripotent stem cells (hESCs) which exhibit an aerobic glycolytic metabolic preference, a metabolic transition not far from the naïve-to-primed bivalency to glycolytic transition of mESCs-to-mEpiSCs (5). Under differentiation conditions, naïve mESCs with a Pkm2 allele knock-in resisted differentiation, whereas a Pkm1 knock-in did not (31). Additionally, the Pkm2 allele knock-in enhances methionine metabolism during differentiation suggesting a pro-oxidative role (31). We previously demonstrate that PKM1 and PKM2 protein abundance significantly increases in formative mEpiLCs (42), therefore, knocking down Pkm2 during this transition could destabilize the pro-oxidative controls necessary for the developmental transition through the formative state during pluripotency transitioning.

Despite employing an isoform targeted approach of the specific splice sites between PKM1 and PKM2, the results of the protein abundance study demonstrate that one of the constructs downregulated PKM2 with an upregulation to PKM1. This was unexpected but is potentially related to the morpholino in question binding across splice regulatory proteins. In this case, the alternative splicing mechanism of PKM1/2 could be impacted by altering splice suppressor and enhancer proteins binding to pre-mRNAs. This would cause a feedback mechanism leading to unintended splicing edits. This alteration has been previously demonstrated in a study examining PKM2 and myotonic dystrophy (43). Future experiments will need to define this possible mechanism. Despite this unexpected effect of PKM MO2, once we consistently detected its disparate effects on PKM2 and PKM1, we realized this experimental approach provided us with the unique opportunity to simultaneously assess the effects of lowering PKM2 and increasing PKM1 on stem pluripotency employing a single morpholino. Our outcomes derived from the use of this morpholino cannot be determined to be exclusively due to either downregulated PKM2 or upregulated PKM1, but uniquely demonstrate that the alteration of expression of these isoforms certainly does dysregulate cell pluripotency marker expression and metabolic enzyme transcript levels. The lack of PKM1 specificity introduced us to an unexpected beneficial strategy that being that our Pkm1/2 morpholinos can induce both downregulation of PKM2 alone and downregulation of PKM2 while simultaneously upregulating PKM1. This occurrence has been demonstrated previously using morpholinos on the alternative spliced gene Proteolipid protein 1 (Plp1/DM20), where Plp1 shift to DM20 alternative splicing (44). The cause for the lack of specificity in the targeted morpholino approach could be due to a duplication mutation as the exclusive exons share a very similar sequence, downstream mutations could further change the exonic structure. A PKM1 specific morpholino is necessary to delineate the role of the M1 isoform in maintaining individual pluripotent states and pluripotent transitioning. A CRISPR or TALEN strategy is possible for an alternatively spliced isoform such as PKM1/2, however, PKM1/2 specificity has been shown using shRNA, germline loss of function, and lentiviral allele-knock in (30,31,39).

Between either end of the pluripotent spectrum exists a recently described and poorly understood executive, formative stage, of which, metabolic trends and PKM expression have yet to be fully delineated (7). Our results demonstrate that altering PKM isoform levels influences metabolic and pluripotent cell surface marker expression. Contrary to the hypothesized direction that transitioning formative and primed-like mEpiLCs would take, knocking down PKM1/2 promoted primed pluripotency associated cell surface marker expression and promoted an enhanced population of cells expressing both high levels of the naïve cell surface marker SSEA1 and the primed cell surface marker CD24. Notably, there was no significant differences in pluripotency associated transcripts following the addition of morpholinos from their non-transfected control states. While previous research has found Pkm2 to play an important role in naïve pluripotency maintenance and reprogramming from somatic cells to either naïve or primed states, these results demonstrate that Pkm1/2 modifications do alter the pluripotent phenotype in transitioning formative and primed-like pluripotent states towards a primed state. The downregulation of PKM2 and upregulation of PKM1 promoted formative state cell expression, however, at this time little is known regarding formative state metabolic preferences (45). Profiling true, stable formative state cells would confirm this potential cell surface marker trend and the role of PKM1/2 during transitioning. Our investigation has shed some light on the metabolic preference of formative state cells through our transcript abundance study. We demonstrate a downregulation of OXPHOS transcripts such as Idh2 and an increase in glycolytic transcripts such as Ldha, these trends suggest the initiation of the aerobic glycolysis and reflect the *in vivo* correlate following post-implantation of the blastocyst (46). Of note, the transcript abundance study found key differences between the formative mEpiLCs and primed mEpiSCs, including an increase in OXPHOS transcripts in Mdh2 and Suclg1 for primed cells. These metabolic markers regulate tumor growth and are players in aerobic glycolysis, suggesting that the formative state has not fully adhered to aerobic glycolysis (47). As metabolically bivalent cells, naïve mESCs transitioning to the formative state rewire their transcriptional and epigenetic landscape, gaining the competency to differentiate into other cell types (7,48). Previous studies demonstrate that reactive oxygen species (ROS)-mediated interactions with mitochondria and nuclear functions are clearly implicated in stem cell fate and potency (49). As PKM1 plays a critical role in the metabolic shunting of pyruvate towards an OXPHOS fate as acetyl-CoA in the mitochondria, the ROS generated by increased PKM1 may promote metabolic reprogramming by inducing a shift to OXPHOS, or an OXPHOS-burst, to increase ROS and stabilize hypoxia inducible factor 1*α*(HIF-1*α*) (50). Such an event could be verified by examining extracellular acidification rate and oxygen consumption rate. The master metabolic regulator HIF-1*α*is activated during instances of hypoxia, decreased ROS and the glycolytic shift towards primed pluripotency. PKM2 can interact with HIF1*α*to further promote aerobic glycolysis (32). Morpholino-induced PKM2 reduction and PKM1 upregulation, suggest that PKM2 reduction promoted the primed CD24 high cell surface marker population when generating formative state mEpiLCs and may conversely indicate a role for PKM1 in promoting naïve pluripotency.

Upregulation of PKM1 appears to have blunted the influence of PKM2 reduction, potentially as a compensatory mechanism. This study demonstrates that formative and primed-like pluripotent states can be effectively distinguished from the naïve, ground, and primed pluripotent states using flow cytometry for the cell surface markers SSEA1 and CD24. Previously, only naïve and primed states have been examined using these markers and here we demonstrate that unique expression dynamics could additionally be utilized for fluorescently activated cell sorting for downstream studies and population purifications (45). This utility could determine transitioning and differentiation efficiencies within the pluripotent continuum and exit during cell lineage specification. This method could be used to help study primordial germ cell-like cell generation along with somatic lineage competency during pluripotent development.

The results of the transcript abundance demonstrate that the glycolysis genes Pgam1 and Gpi significantly decreased in primed mEpiSCs following PKM2 downregulation. This is not surprising as Pkm2, Pgam1, and Gpi are heavily implicated in aerobic glycoloysis and biosynthesis. Downregulating a key protein such as PKM2 appears to have downstream effects on other aerobic glycolysis-associated genes and may disrupt aerobic glycolysis in cells that have achieved true primed pluripotency (51,52). Importantly, the genes Hk2 and Eno1 significantly increased in primed mEpiSCs, and Hk2 also significantly increased in primed-like mEpiLCs following PKM2 downregulation and PKM1 upregulation. This follows the currently described preferences for primed pluripotency, and exit of the naïve state, by promoting a glycolytic preference over OXPHOS, and elevated aerobic glycolysis transcription of these two critical genes (53). The transcript abundance results also demonstrate that the OXPHOS genes Mdh2, Fh1, and Suclg1 all significantly increased when PKM2 was downregulated and PKM1 was upregulated in primed-like mEpiLCs. This would further promote the primed pluripotent state as Mdh2 is implicated in feeding aerobic glycolysis through nicotinamide adenine dinucleotide regeneration (NAD), thus supporting the glycolytic shift (54). Fumarate hydratase (Fh1) processes the accumulated fumarate to activate a hypoxia response (55,56). The increased PKM1 may contribute towards compensating towards fumarate accumulation, however, increased Fh1 relative to the control primed-like cells and even the naïve mESCs is unexpected and may play a new role in generating the unique SSEA1 high, CD24 high cell population. Suclg1 works to generate ADP and succinyl-CoA in the tricarboxylic acid cycle and can promote substrate level phosphorylation even in the absence of oxygen, and thus can work within the shift towards aerobic glycolysis model in primed-like mEpiLCs (57). Metabolic profiling at the protein level through immunoblots and non-denaturing gels paired with live cell acute measure of extracellular acidification rate and oxygen consumption rate will help elucidate metabolic trends in the formative state and PSCs treated with PKM morpholinos. As both the downregulated PKM2 and the combination of downregulation of PKM2 and upregulation of PKM1 yielded similar levels (mean of 36.4% and 38.7% respectively) of unique population of SSEA1 high and CD24 high primed-like mEpiLCs, the effects of PKM1 upregulation are either not strong enough to compensate or do not have a role in the transition out of the formative state to primed state pluripotency. This trend could additionally be in response to, or in addition to the significant increase in the ratio of PKM1/PKM2 with the PKM MO 2 treatment. The metabolic transcript abundance results demonstrate modulating PKM1 and PKM2 expression appears to impact the primed-like mEpiSCs and primed mEpiSCs. This may promote the unique population of naïve and primed cell surface marker expressing cells in the primed-like mEpiSCs, and likely disrupts primed pluripotency in mEpiSCs by displacing aerobic glycolysis (Fig. 8.). Our strategy of targeting PKM1/2 splice events resulted in an elevated phosphorylated-PKM2 (pPKM2) to total- PKM2 ratio showing the cumulative expression of both homo-tetrameric and dimeric (pPKM2) conformations. This result implicates pPKM2 as playing a role in generating this novel SSEA1 high and CD24 high expression pattern following the transition from the formative state to a primed-like pluripotency. Previously, we demonstrated maintenance of the ratio of PKM1/PKM2 protein abundance throughout the pluripotent continuum in murine cells (42), by modulating the ratio of pPKM2/PKM2 with morpholinos, a further promotion of glycolysis and enhanced metabolic shifting from bivalency to aerobic glycolysis is possible.

Here, we demonstrate that employment of upregulation of PKM1 with downregulation of PKM2 promoted primed pluripotent stem cell populations (CD24 High) when generating formative state mEpiLCs. Importantly, downregulating PKM2 with PKM1 upregulation does result in a significant increase in CD24 High expressing cells. In this instance, the pro-OXPHOS nature of Pkm1 may promote the bivalent nature of mESCs to counter the pro-transitioning influence of downregulating PKM2. Through germline deletion of PKM2 in mice, it was found that PKM1 becomes the predominant isoform in all cells of the developing PKM2-null mouse compensating for the loss of PKM2, the traditional predominant isoform during development (39). These results suggest that downregulating PKM2 appears to yield a population expressing both naïve and primed cell surface markers. Interestingly, another intermediate pluripotent cell state referred to as ‘f-class’ cells have been shown to contain a similar population of SSEA1+/CD24+ cells (58,59). These f-class cells are generated in rare populations following extended transgene expression of reprogramming factors Oct4, Klf4. Sox2 and c-Myc (58). When the formative pluripotent state was hypothesized, the notion that formative cells could occupy the transcriptional profile of both naïve and primed states was suggested, potentially disrupting the fine-tuning pro-oxidative controls associated with the Pkm2 isoform during a transition that promotes the formative state *in vitro* (7). When taken together, this study promotes PKM1 and PKM2 having a role in aerobic glycolysis as upregulation in formative and primed-like mEpiLCs and mEpiSCs promotes glycolytic genes and primed pluripotency associated CD24 expression. This is surprising as PKM2 is associated with aerobic glycolysis, and PKM1 has only recently been implicated in specific cancers (60,61). These findings promote Pkm1 and Pkm2 having distinct roles in metabolism and pluripotency (26,42,61) along with contributing to the growing body of evidence that metabolism is a driver of pluripotency state.

## Acknowledgements

Flow cytometry was completed at the London Regional Flow Cytometry Facility at Robarts Research Institute of The University of Western Ontario, London, Ontario, Canada. Pluripotent cell lines were generously gifted from Dr. Janet Rossant from The Hospital for Sick Children, Toronto, Ontario, Canada. Morpholinos were designed by Dr. Jon Moulton of Gene Tools LLC, Philomath, Oregon, USA. This research was funded by a Canadian Institutes of Health Research operating grant to A.J.W. and D.H.B. and Natural Sciences and Engineering Research Council of Canada grant to D.H.B. The funders had no role in study design, data collection and analysis, decision to publish, or preparation of the manuscript.

## CRediT Author Statement

**Joshua G. Dierolf:** Conceptualization, Methodology, Validation, Formal Analysis, Investigation, Writing – Original Draft, Writing – Review & Editing, Visualization **Hailey Hunter:** Methodology, Writing – Review & Editing **Dean H. Betts:** Resources, Writing – Review & Editing, Funding Acquisition, Supervision **Andrew J. Watson:** Writing – Review & Editing, Funding Acquisition, Supervision

## Additional information

The authors have no competing interests to disclose.

---

**Supplemental FIG. 1.**
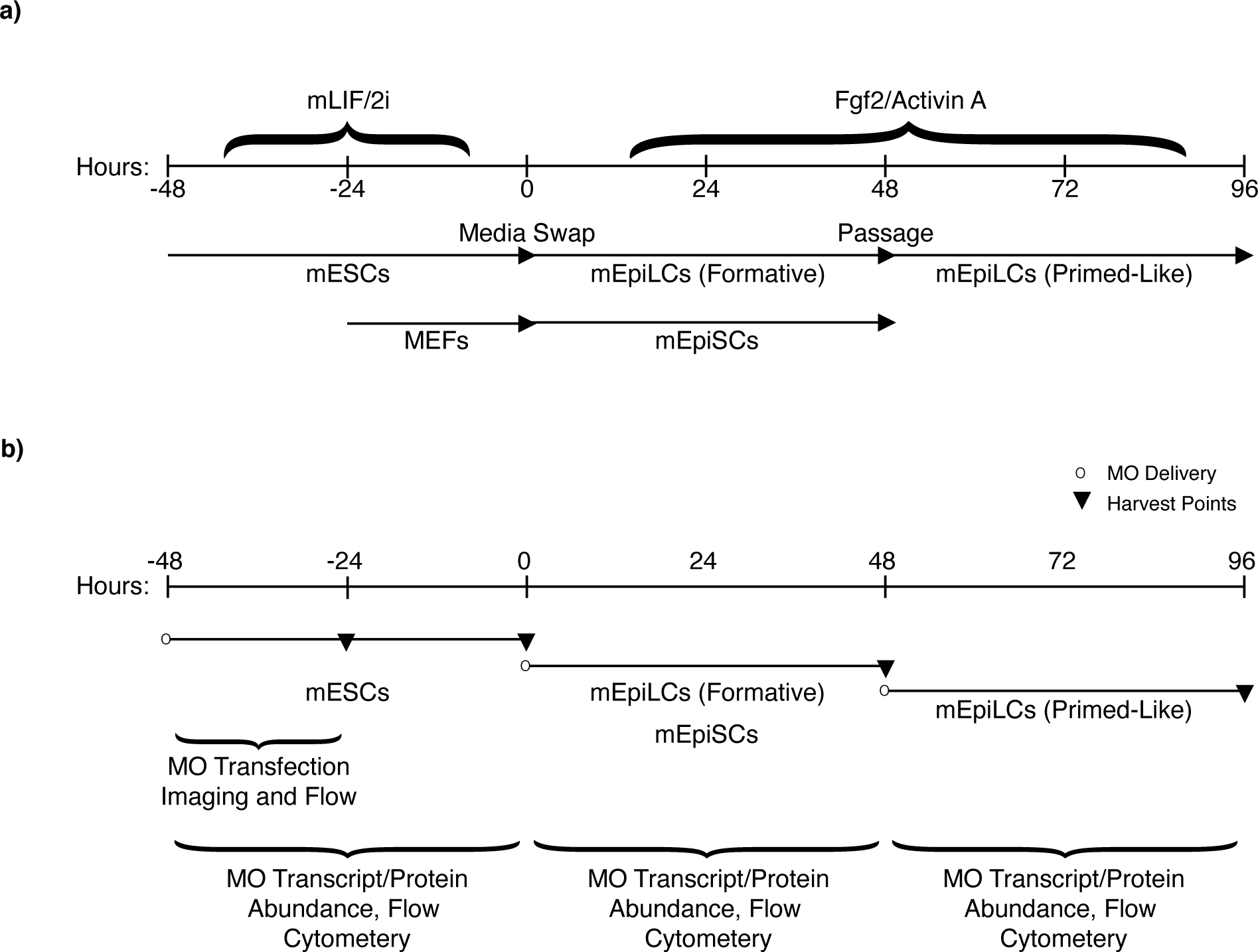
mESC, mEpiLC, and mEpiSC culture and timing schematic. Ground state, naïve mESCs, formative/primed-like transitioning to mEpiLCs, and primed mEpiSCs experimental planning schematic. a) Experimental plating set-up and media transitioning from mESCs into formative and primed-like mEpiLCs through media supplementation and timing. b) Morpholino scrape delivery and incubation timelines described per cell type in each experimental study component.

